# Risk of differential cancer types over age in families with Li-Fraumeni syndrome: a validation study using multi-center cohorts

**DOI:** 10.1101/567727

**Authors:** Seung Jun Shin, Elissa Dodd, Gang Peng, Jasmina Bojadzieva, Jingxiao Chen, Chris Amos, Phuong L. Mai, Sharon A. Savage, Mandy L. Ballinger, David M. Thomas, Ying Yuan, Louise C. Strong, Wenyi Wang

## Abstract

Li-Fraumeni syndrome (LFS) is a rare hereditary cancer syndrome associated with an autosomal dominant mutation inheritance in the *TP53* tumor suppressor gene and a wide spectrum of cancer diagnoses. An accurate estimation of the penetrance of different cancer types in LFS is crucial to improve the clinical characterization and management of high-risk individuals of LFS. Our competing risk-based statistical model incorporates the pedigree structure efficiently into the penetrance estimation and corrects for the ascertainment bias. A set of *TP53* penetrance for three cancer types (breast, BR/sarcoma, SA/others, OT) involved in LFS is then estimated from 186 pediatric-sarcoma families collected at MD Anderson Cancer Center. The penetrance was then validated on a mixed cohort of clinically ascertained families with cancer (total number of families=668). The age-dependent onset probability distributions of specific cancer types are different. For breast cancer, the *TP53* penetrance goes up at an earlier age than the reported *BRCA1/2* penetrance. We validated the prediction performance of the penetrance estimates via two independent cohorts combined (BR=85, SA=540, OT=158). We obtained areas under the ROC curves (AUCs) of 0.92 (BR), 0.75 (SA), and 0.81 (OT). Our R package, LFSPRO, provides risk estimates for the diagnosis of breast cancer, sarcoma, other cancers or death before cancer diagnosis in future.

**Significance:** Cancer-specific penetrance can facilitate clinical characterization of LFS and will contribute to the management of families at high risk of LFS.

## INTRODUCTION

Li-Fraumeni syndrome (LFS) is a rare familial cancer syndrome that is characterized by early cancer onset and a wide diversity of tumor types in contrast to other cancer syndromes^1,2^. The initial identification of the syndrome was through aggregated family cancer history with the major hallmarks including bone and soft tissue sarcoma, breast cancer, adrenal cortical cancers, brain tumors, and leukemia^3–7^, but recent studies have shown that its spectrum gets more diverse and includes lung cancer^7^ and prostate cancer^8^.

LFS is known to be associated with deleterious germ-line mutations in the *TP53* tumor suppressor gene. It is crucial to accurately estimate the penetrance of *TP53* mutations in order to provide better clinical management to the individuals at high risk of LFS. Due to the wide spectrum of LFS cancers, it is likely that specific *TP53* mutations have different effects for different cancer types. However, in recent literature^9^, the cancer type information is collapsed and the disease status is simply dichotomized (cancer or not). This type of penetrance, which we call overall-cancer penetrance, ignores the cancer type information and fails to quantify the cancer-specific effect of a *TP53* germline mutation. It is necessary to estimate a cancer-specific penetrance in order to not only reveal the cancer-specific effect of the *TP53* mutations but also improve the risk prediction performance by accommodating more detailed disease histories. However, it is a complex task to take into account multiple types of cancers simultaneously since onsets of different cancers are competing with others.

Recently, a completing risk model was developed that captures the nature of the multiple cancers associated with LFS^10^. In this article, we provide a set of cancer-specific age-at-onset penetrances of *TP53* under the model proposed^10^. First, we provide a more realistic penetrance by taking into account deaths unrelated to LFS as a baseline competing risk factor. Second, we utilize the pedigree information into the estimation which can substantially improve estimation accuracy and efficiency by including more patient data where genotype information is unknown^9^. Recently cancer-specific observations of LFS have been reported^11^, however, they calculate the cumulative incidence for different cancer types based on deleterious *TP53*+ individuals only. Third, we carefully handle the ascertainment bias inevitable in rare disease studies like LFS and provide an unbiased penetrance estimate that can be applied to the general population.

As mentioned above, an accurate cancer specific characterization of LFS is an extremely difficult task due to its nature involving such a wide spectrum of cancer diagnoses. We hereby provide a new set of cancer-specific penetrance estimates for individuals with different *TP53* mutation status, sex and age. Cancer-specific penetrance estimates could be used by genetic counselor and physicians to provide more detailed counseling to families that have newly been found to harbor a *TP53* germline mutation.

## METHODS

### Cancer-specific penetrance

Due to the rare prevalence of most cancer types in LFS, we grouped all cancers related to LFS into three types: breast cancer, sarcoma, and all others. In our competing risk model, each different cancer type is competing with all others as the first event. As a baseline competing event, we consider deaths irrelevant to LFS. For example, breast cancer is observed only if the breast cancer appears before other cancers or death. Therefore, we have four competing events: 1) breast cancer, 2) sarcoma, 3) other cancers and 4) dying without having been diagnosed with cancer.

The age-at-onset cancer-specific penetrance is defined by a cumulative probability (up to a certain age) of experiencing a particular competing event prior to all others, otherwise referred to as cumulative incidence in the competing risk literature. This is different from a net probability of being diagnosed as a particular cancer type when possible competing risks are removed, which consequently overestimates the actual risk. Considering the sex (male and female) as an additional non-genetic covariate, we estimate a complete set of penetrance estimates for the four competing events at four different configurations of covariates, i.e., {male, female} * {wildtype, mutated}. Introducing death events as a baseline competing risk, we can estimate the penetrance for death irrelevant to LFS, which is a natural occurrence and a crucial factor for assessing real cancer risk. The overall cancer penetrance without considering its subtype can be obtained by simply accumulating the cancer-specific penetrance. Similarly, the *TP53* mutation penetrance for either experiencing any cancer diagnosis or dying without cancer is obtained by summing up the overall cancer penetrance plus the death penetrance. The penetrance estimates for the wildtype population are directly comparable to the corresponding Surveillance, Epidemiology, and End Results (SEER) estimates without any further modification (**Figure 1F**).

**Figure 1.**
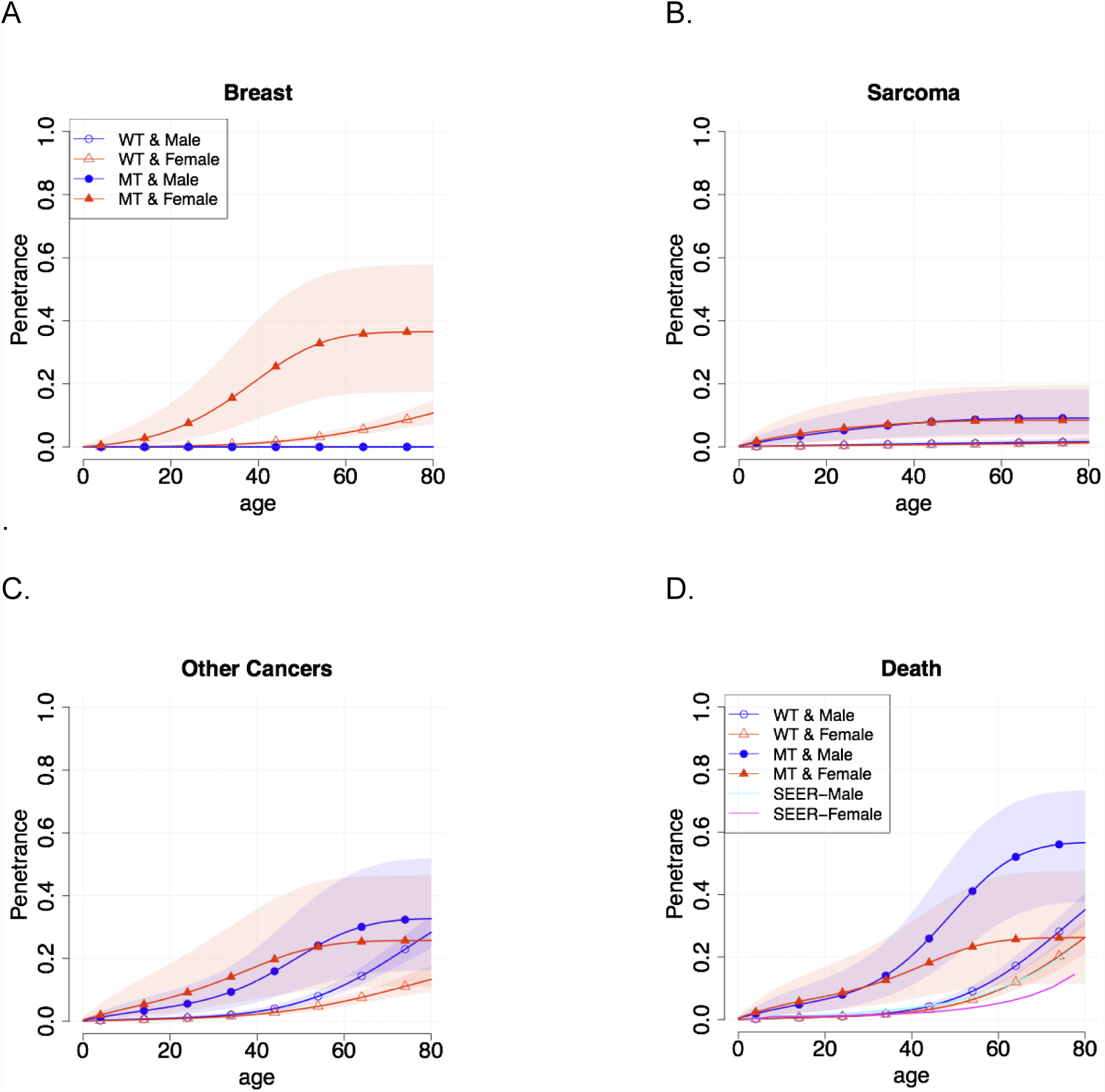
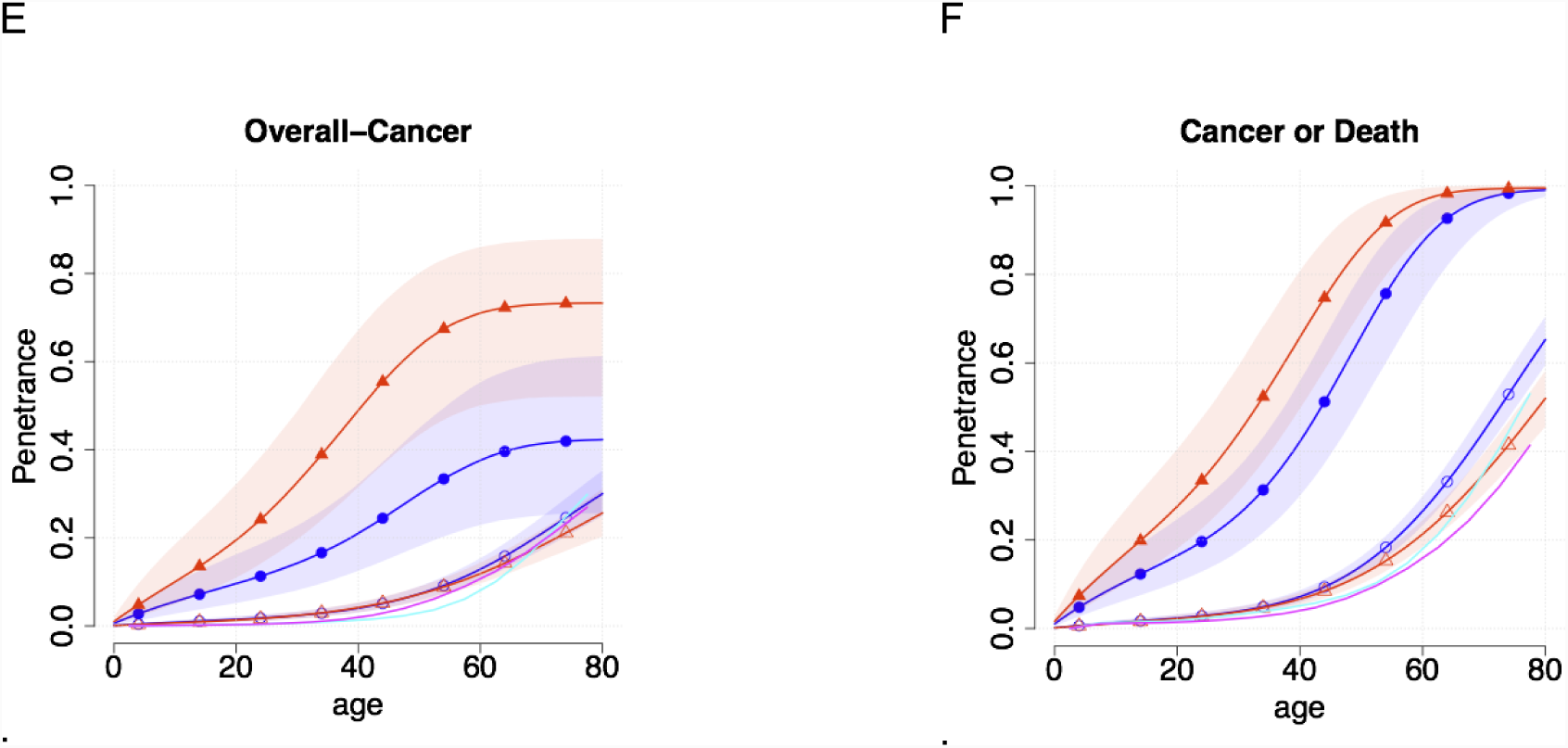
Cancer-specific penetrance estimates for *TP53* mutation carriers and noncarriers from the pediatric sarcoma cohort. Three cancer-specific penetrances (A-C) estimated from the pediatric sarcoma cohort are depicted along with the corresponding 95% confidence bands. As byproducts death penetrance (D), overall-cancer penetrance (E), and cumulative probability of either having any cancers or dying without cancer (F) are given. Their non-carrier penetrances are compared to the corresponding SEER estimates.

### Model and Estimation

Cancer-specific hazard is considered under a competing risk framework. In order to incorporate pedigree structure and family cancer history into the estimation, familywise likelihood^9^ and family-specific random frailty are introduced. Ascertainment bias is adjusted to obtain unbiased penetrance estimates^12^. Finally, we refer to our methods paper^10^ for complete details about the model and the associated Bayesian estimation scheme via Markov chain Monte Carlo (MCMC) algorithm.

### Study Population

We used a cohort of 186 families collected via patients (probands) with childhood soft-tissue sarcoma or osteosarcoma treated at The University of Texas M.D. Anderson Cancer Center (MDACC, Houston, TX) from 1944 to 1983 to train the model. Details of the data have been published elsewhere^7,13–16^. Medical records and death certificates were used to confirm cancer histories, where possible. We define the subjects as affected only if diagnosed with malignant tumors. Cancers are classified by three categories: breast cancer, sarcoma (osteosarcomas and soft tissue sarcomas), and other cancer (brain, adrenal, lung, leukemia and all other cancers). Both linkage and segregation analysis were carried out for selecting high-risk families prior to testing *TP53*. After a *TP53* mutation is identified for the proband, *TP53* mutation status was determined by PCR sequencing of exons 2-11. If a *TP53* mutation was identified, all first-degree relatives of the proband and any other family member with an increased risk of carrying the mutation were tested. Extending germline testing to additional family members based on mutation status instead of phenotypes should not introduce ascertainment bias during analysis^7,9^. Individuals unavailable for testing who are linked to or between confirmed mutation-carriers were considered inferred mutation carriers. Immediate family members were not tested if the proband did not have *TP53* mutation. Out of 186 families, 17 are referred to as (*TP53* mutation) positive families, which have at least one mutation carrier identified, and the remaining 182 families, in which there is no carrier identified, are considered wild-type. See **Table 1** for detailed summary of the MDACC data.

**Table 1.** MDACC Cohort Summary: average ages of diagnosis or at death are given in parentheses.

| Gender | Genotype | Breast Cancer |  | Sarcoma |  | Other-Cancers |  | Death |  | Lost Follow-up |  | Total |  |
| --- | --- | --- | --- | --- | --- | --- | --- | --- | --- | --- | --- | --- | --- |
| Male | Unknown | 0 | (NA) | 11 | (21.3) | 130 | (55.3) | 232 | (55.7) | 1043 | (39.8) | 1416 | (43.9) |
|  | Wildtype | 0 | (NA) | 76 | (9.6) | 30 | (57.1) | 38 | (63.6) | 257 | (40.0) | 401 | (37.9) |
|  | Mutation | 0 | (NA) | 12 | (13.2) | 27 | (45.3) | 1 | (47.0) | 8 | (32.5) | 48 | (35.2) |
|  | Subtotal | 0 | (NA) | 99 | (11.3) | 187 | (54.2) | 271 | (56.7) | 1308 | (39.8) | 1865 | (42.3) |
| Female | Unknown | 39 | (55.6) | 4 | (8.2) | 62 | (46.5) | 168 | (59.7) | 1035 | (41.5) | 1308 | (44.5) |
|  | Wildtype | 8 | (53.2) | 96 | (11.1) | 17 | (54.5) | 45 | (64.4) | 297 | (41.2) | 463 | (38.0) |
|  | Mutation | 19 | (33.2) | 12 | (9.2) | 7 | (23.1) | 1 | (22.0) | 8 | (37.9) | 47 | (25.8) |
|  | Subtotal | 66 | (49.0) | 112 | (10.8) | 86 | (46.0) | 214 | (60.5) | 1340 | (41.4) | 1818 | (42.3) |
| Total |  | 66 | (49.0) | 211 | (11.1) | 273 | (51.5) | 485 | (58.1) | 2648 | (40.6) | 3683 | (42.3) |

### Validation Study

We validated the penetrance estimates by comparing disease statuses and cancer-specific risk of genotype-observed subjects in the MDACC data. The cancer-specific risk for a genotype-observed individual is readily obtained by differentiating the estimated cancer-specific penetrance estimates. However, there are two potential problems in this validation study. First, large proportions of genotype-observed individuals are probands, who have excessively high risk for a *TP53* mutation. We exclude probands in the validation study in order to mitigate this bias. Second, we use the MDACC data for both estimating and validating penetrances, and consequently validation results become too optimistic. To tackle this issue, we conducted validation analysis for two different prospective cohorts. One cohort is recruited to the International Sarcoma Kindred Study (ISKS) through adult-onset sarcoma patients from six clinics in Australia. Cancers were verified by Australian and New Zealand cancer registries and death certificates^17^. The proband’s *TP53* mutation status was determined by PCR sequencing, high-resolution melt analysis and multiplex ligation-dependent probe amplification to detect large deletions or genomic rearrangements^17^. The ISKS cohort consist of 582 families out of which nineteen are *TP53* mutation positive families (**Table 2**). The second independent cohort was recruited to the National Cancer Institute (NCI). The NCI LFS study is a long-term prospective, natural history cohort study that started in August 2011, and includes individuals meeting the classic LFS^18^ or Birch’s LFL^4^ criteria, having a pathogenic germline *TP53* mutation or a first- or second-degree relative with a mutation, or having a personal history of choroid plexus carcinoma, adrenocortical carcinoma, or at least 3 primary cancers (NCT01443468). A detailed family history questionnaire was obtained, including information on birth date, vital status, date or age at death if deceased, history of cancer, and if positive, type and year of diagnosis or age at diagnosis, for all first-, second-, and third-degree relatives and any other extended family members for whom the information was available. There were 2,676 individuals from 102 families included in this NCI LFS cohort^19^ (**Table 3**).

**Table 2.** ISKS Cohort Summary

| Gender | Genotype | Breast Cancer |  | Sarcoma |  | Other-Cancers |  | Death |  | Lost Follow-up |  | Total |  |
| --- | --- | --- | --- | --- | --- | --- | --- | --- | --- | --- | --- | --- | --- |
| Male | Unknown | 2 | (53.5) | 12 | (29.2) | 696 | (60.6) | 2282 | (64.5) | 5586 | (42.4) | 8578 | (49.7) |
|  | Wildtype | 0 | (NA) | 265 | (45.6) | 47 | (50.3) | 0 | (NA) | 6 | (54.2) | 318 | (46.5) |
|  | Mutation | 0 | (NA) | 9 | (28.3) | 4 | (33.2) | 0 | (NA) | 1 | (22.0) | 14 | (29.3) |
|  | Subtotal | 2 | (53.5) | 286 | (44.4) | 747 | (59.8) | 2282 | (64.5) | 5593 | (42.4) | 8910 | (49.6) |
| Female | Unknown | 243 | (55.2) | 18 | (38.3) | 544 | (56.6) | 1857 | (68.4) | 5360 | (43.8) | 8022 | (50.7) |
|  | Wildtype | 15 | (52.9) | 212 | (45.9) | 38 | (41.8) | 0 | (NA) | 11 | (36.5) | 276 | (45.3) |
|  | Mutation | 3 | (32.0) | 6 | (43.3) | 1 | (10.0) | 0 | (NA) | 3 | (31.0) | 13 | (35.3) |
|  | Subtotal | 261 | (54.8) | 236 | (45.2) | 583 | (55.5) | 1857 | (68.4) | 5374 | (43.8) | 8311 | (50.5) |
| Total |  | 263 | (54.8) | 522 | (44.8) | 1330 | (57.9) | 4139 | (66.2) | 10967 | (43.1) | 17221 | (50.0) |

**Table 3.** NCI Cohort Summary

| Gender | Genotype | Breast Cancer |  | Sarcoma |  | Other-Cancers |  | Death |  | Lost Follow-up |  | Total |  |
| --- | --- | --- | --- | --- | --- | --- | --- | --- | --- | --- | --- | --- | --- |
| Male | Unknown | 1.0 | (60.0) | 27 | (25) | 238 | (46.8) | 0 | (NA) | 876 | (25.2) | 1142.0 | (29.7) |
|  | Wildtype | 0 | (NA) | 3 | (56.7) | 4 | (39.5) | 0 | (NA) | 107 | (22.4) | 114.0 | (23.9) |
|  | Mutation | 0.0 | (NA) | 18 | (38.7) | 29 | (43) | 0 | (NA) | 31 | (27.5) | 78.0 | (35.8) |
|  | Subtotal | 1.0 | (60) | 48.0 | (32.1) | 271. | (46.3) | 0 | (NA) | 1014.0 | (24.9) | 1334.0 | (29.6) |
| Female | Unknown | 143. | (50.7) | 22 | (30.1) | 151. | (42.4) | 0 | (NA) | 756 | (25.6) | 1072.0 | (31.4) |
|  | Wildtype | 16.0 | (54.3) | 3 | (41.3) | 8.0 | (49.8) | 0 | (NA) | 108 | (23.4) | 135.0 | (29) |
|  | Mutation | 51.0 | (43.1) | 24 | (35.6) | 27.0 | (35.9) | 0 | (NA) | 33 | (20) | 135.0 | (34.7) |
|  | Subtotal | 210. | (49.1) | 49.0 | (33.5) | 186. | (41.8) | 0 | (NA) | 897.0 | (25.1) | 1342.0 | (31.5) |
| Total |  | 211 | (49.2) | 97 | (32.8) | 457 | (44.5) | 0 | (NA) | 1911 | (25) | 2676 | (30.5) |

### Prevalence and de novo Mutation Rate

We assume the *TP53* mutation follows Hardy-Weinberg equilibrium. The mutation prevalence is 0.0006^19^. The frequencies for the three genotypes (homozygous reference, heterozygous and homozygous variant) are 0.9988004, 0.00059964 and 3.6e-07, respectively. We used 0.00012 as a default value of *de novo* mutation rate when evaluating the familywise likelihood^19,20^.

### Validation study design

We evaluated the model prediction performance on cancer-specific risk using the average annual risk computed with our cancer-specific *TP53* penetrance estimates. The cancer-specific risk was calculated as the cumulative probability of developing one type of cancer divided by the follow-up time. The receiver operating characteristic (ROC) curve was used to evaluate the sensitivity and specificity of predicting incidence of a specific cancer type, using the estimated risk probability at various cutoffs. We also provide 95% bootstrap confidence intervals for AUC estimates^21,22^.

## RESULTS

### Cancer Specific Penetrance Estimates

As shown in **Table 1**, the pediatric sarcoma cohort at MDACC provided clinical outcomes of 3,683 individuals from 168 families. Among them, there are 66 breast cancer cases, 211 sarcoma cases, 273 other cancers, 485 deaths and 2,648 lost to follow-up. Within each of the four competing events, 27 breast cancer cases (41%), 196 sarcoma cases (93% - explained by ascertainment as most of them are probands), 57 other cancers (21%) and 85 deaths (18%) had known *TP53* genotype status. Our competing risk and pedigree-based hazard model accounts for outcomes of individuals without genotype status by using Mendelian inheritance that assigns these individuals with weights for being a carrier or a non-carrier. This approach substantially increased the sample size and the accuracy for penetrance estimation.

The set of cancer-specific penetrances estimated from the pediatric sarcoma cohort are depicted in **Figure 1**. **Figures 1A**, **1B**, and **1C** show the three cancer-specific penetrance estimates for breast cancer, sarcoma, and other cancers combined, respectively. Note that the estimated cancer-specific penetrance estimates display completely different patterns for different cancers, which could be identified only through the proposed cancer-specific approach. **Figure 1D** depicts the penetrance for death before cancer diagnosis. **Figure 1E** and **1F** are the combined penetrance of overall cancer and cancer or death, respectively. In each panel, we present four penetrance estimates each of which corresponds to one of configurations of sex and genotype.

Risk of breast cancer greatly increases between 20 and 40 years of age for female *TP53* mutation carriers (**Figure 1A**). We further compare it to the female penetrance estimate of *BRCA1/2* which are two well-known breast cancer susceptibility genes (**Figure 2**). Since *BRCA1/2* penetrance ignores possible competing events such as death and other cancers they are not directly comparable to our *TP53* breast cancer penetrance. However, our estimates can be easily converted to the corresponding net counterpart that ignores competing risks^17^. Through the head-to-head comparison, we observed that the female *TP53* mutation carrier has excessively high probability of developing breast cancer before 25, which is not the case for *BRCA*1/2 (**Figure 2**). Therefore, we provide quantitative evidence for a very early-onset breast cancer being regarded as a clinical evidence of mutations in *TP53*. In addition, the noncarrier penetrance of both *TP53* and *BRCA*1/2 do coincide, which supports the validity of our estimates (**Figure 2**).

**Figure 2.**
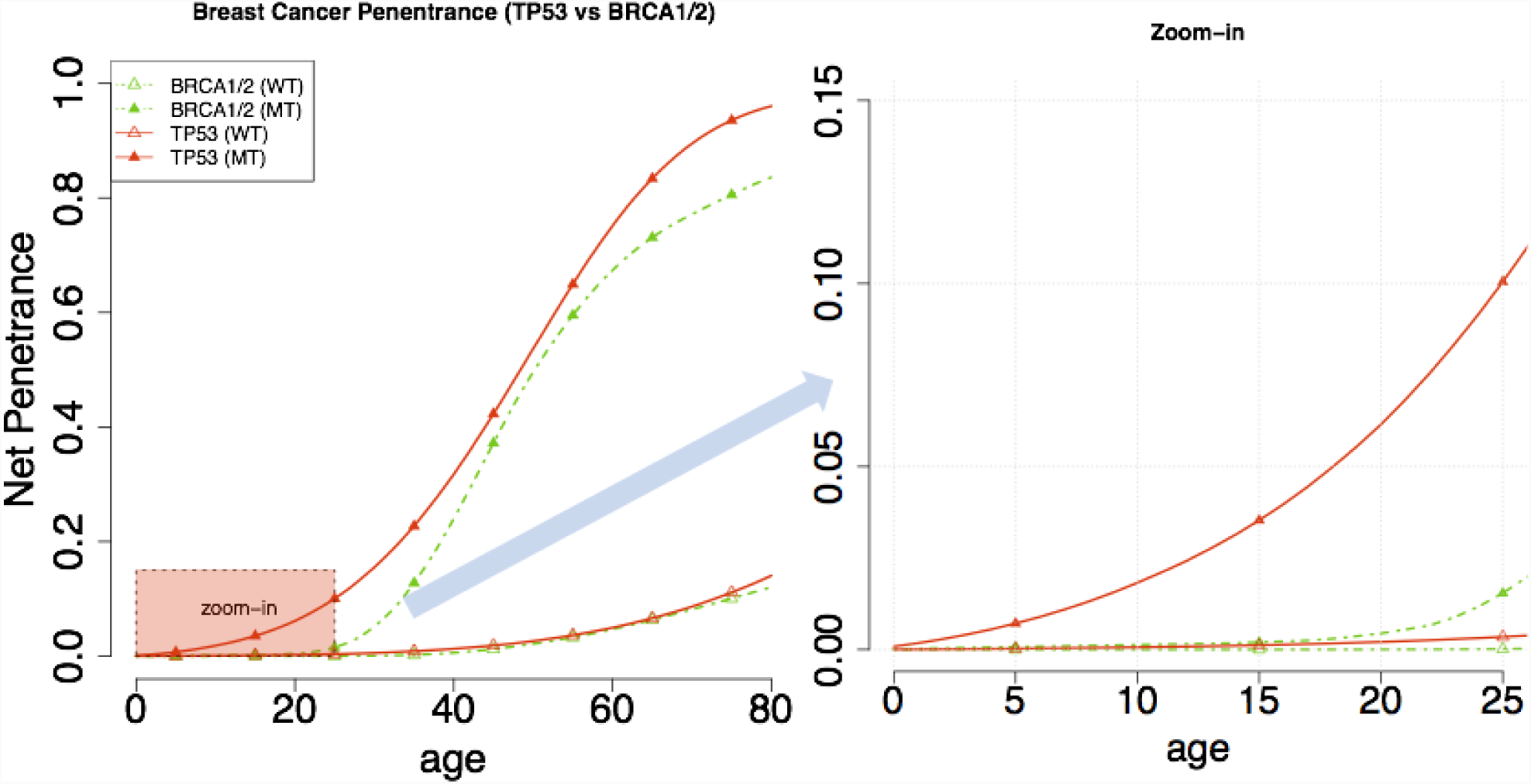
Comparison of TP53 net penetrance to the BRCA1/2 net penetrance in breast cancer. A female *TP53* mutation carrier has excessively high probability of the breast cancer in early age, which is not the case for *BRCA*1/2. A very early-onset breast cancer can be regarded as a clinical evidence of mutations in *TP53*. It is observed that non-carrier penetrances of both *TP53* and BRCA1/2 coincide.

The sarcoma specific *TP53* penetrance increases by about 10% from wild-type individuals from birth to 45 years of age. Although the MDACC pediatric sarcoma cohort is not a random sample and contains increased number of cases of sarcoma compared to the general population, we have the sarcoma penetrance estimated at very low risk for wild-type individuals (**Figure 1B**). This demonstrates our estimates successively corrected for the ascertainment. Recalling that the penetrances are estimated from MDACC data collected through pediatric sarcoma patients, it is not surprising to observe the carrier sarcoma penetrance rapidly increases at an early age and then plateau.

The other-cancer-combined penetrance for *TP53* carriers is noticeably different that the wild-type individuals. For females, the risk of other cancer increases from 5% to 22% between 5-55 years of age and then begins to plateau. For males there is a significant penetrance increase between 35 and 65 years of age.

Interestingly, the penetrance of death before cancer for *TP53* carriers was different than that of non-carriers and of the general population SEER estimates which was used for validation purposes. Missing information in cancer outcomes in obligate carriers might contribute to this observation. We also observed higher death rate among the non-carriers (**Figure 1D**). The MDACC pediatric sarcoma cohort family data date back from the 1940’s (**Supplementary Figure 1**) while the most recent SEER estimates are from 2008 to 2010. As shown in **Supplementary Figure 2**, the mortality rate in the US population has decreased over time hence the observed discrepancy.

Next, **Figure 1E** depicts the overall-cancer penetrance and **Figure 1F** shows probabilities of either having any cancer or dying without cancer. Again, we observed high concordances between the non-carrier penetrances to the corresponding SEER estimates for any cancer, and any cancer or dying without cancer.

### Validation Results

**Table 2** and **Table 3** provides summaries of the validation datasets, the ISKS and NCI cohorts. **Figure 3** depicts Receiver Operating Characteristic (ROC) curves of cancer-specific risk estimates and their actual cancer status of genotype-observed subjects. **Figure 3A** are the results for the MDACC data. A total of 772 tested individuals were used, excluding probands. It is not surprising that the cancer-specific risk estimates directly from the cancer-specific penetrances estimated from the MDACC data performed very well with the area under the ROC curve (AUC) equal to 0.855 (95% CI: 0.781, 0.929) for breast cancer, 0.76 (95% CI: 0.589, 0.93) for sarcoma, and 0.749 (95% CI: 0.689, 0.808) for other cancers. **Figure 3B** shows the validation result for the independent cohorts, ISKS and NCI, which are not used for penetrance estimation. A total of 402 tested individuals (38 from ISKS, 364 from NCI) were used, excluding probands. We observed that the penetrance estimates accurately quantifies the associated cancer risks even for these two independent data sets. The AUCs are 0.914 (95% CI: 0.872, 0.957) for breast cancer, 0.748 (95% CI: 0.678, 0.818) for sarcoma, and 0.805 (95% CI: 0.745, 0.865) for other cancers. These results strongly support the validity of the cancer specific estimates.

**Figure 3.**
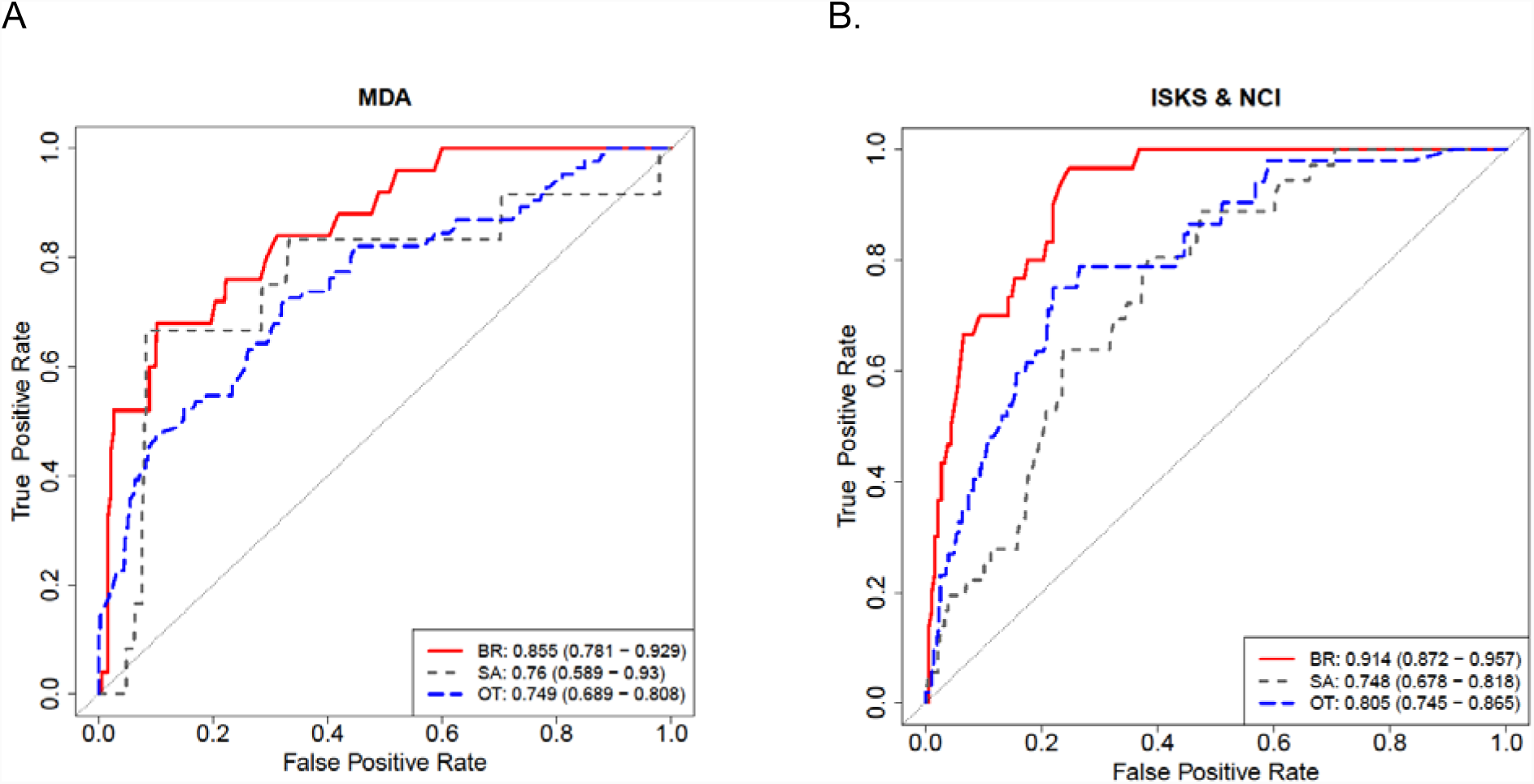
Validation Results using the training dataset (MDA) and the testing dataset (ISKS+NCI). ROC curves that compare cancer-specific risks obtained from the estimated penetrances and disease status of genotypes-observed subjects only are depicted for MDACC data, and the ISKS and NCI data. Probands from all cohorts are excluded to mitigate the bias induced by ascertainment. (BR – breast cancer, SA – sarcoma, OT – other cancer)

## DISCUSSION

We provide the first set of cancer-specific penetrance estimate of *TP53*. These cancer-specific penetrances are modeled under a competing risk framework since all the cancers are competing with others as the first event. A baseline competing event of death irrelevant to the cancers is also incorporated in the estimation. This leads us to precisely estimate a crude risk of cancer without additional calibration based on external data source. Our final penetrance estimates for specific cancer types are age-of-diagnosis-dependent and vary with cancer type, sex and *TP53* mutation status. Based on the new penetrance estimates, we observed the risk for breast cancer is higher than that from *BRCA*1/2 mutations before age 25. Our penetrance estimates are validated using independent cohorts from NCI and ISKS. We have integrated the new penetrance estimates in our risk prediction software LFSPRO and upgraded to version 2.0.0 to provide risk estimates, which is freely available at https://bioinformatics.mdanderson.org/public-software/lfspro/.

Like other cancer syndromes, LFS is extremely rare and it is challenging to collect sufficient amounts of data to be analyzed. However, the pediatric sarcoma cohort from MDACC is a large and comprehensive cohort that only looked at children with a sarcoma diagnosis, excluding any bias of only obtaining families based on LFS clinical criteria^18,23,24^. For validation purposes, we combined the NCI and ISKS cohort for a larger data set and to combine different ascertainment strategies.

With cancer-specific penetrances available, we developed an associated risk assessment extension to the LFS mutation carrier risk prediction tool, LFSPRO R package, based on the BayesMendel model^19,25^. The lfspro.mode function set to “1st.cs” in the updated R package uses an individual’s family history to estimate the risk of breast cancer diagnosis, sarcoma diagnosis, other diagnosis or death for the next 5, 10, 15 and 20 years. Current LFS standard screening protocols^26^ have been instituted in clinics throughout the world^27–31^. At MDACC, the Li-Fraumeni Education and Early Detection (LEAD) Program consists of “a centralized approach to patient management, screening exams performed at the same institution, including whole-body MRI and brain MRI, using same technique and same machines enabling more consistent comparison of findings between and across interval exams in patients and a multidisciplinary team that reviews all findings and patient issues to develop follow up recommendations”^29^. Implementation of cancer-specific penetrance estimation in LFS screening programs could give patients a more complete picture of predicted risk. With a 90% lifetime risk of cancer in *TP53* mutation carriers^7,13^, and a near 100% risk of cancer for female *TP53* mutation carriers by age 70^9^, patients that have tested positive for a germline mutation are constantly on alert for when a cancer diagnosis will occur. Our goal is for LFSPRO is be used as a tool for genetics counselors and clinicians that work with *TP53* mutation carriers to provide more information to their patients during their screening visits. However, this estimation is only for the first primary cancer diagnosis. Unless a mutation has already been identified within a family, most individuals do not find out they are a *TP53* germline mutation carrier until after their first or second primary. Clinical use of the penetrance will be most useful for genetic counselors during mutation testing results disclosure prior to the first cancer diagnosis.

In summary, we provide cancer-specific penetrance estimates of *TP53* mutation carriers with improved resolution that allows us to utilize the cancer type information. Looking deeper into the cancer-specific penetrance of LFS leads to more information for patients and their clinicians. Though treatment targets are being researched, patients long for more information on how LFS affects them now. Adding to the current understanding of the cancer-specific penetrance leads to better understanding of the disease and improve risk management of healthy individuals from families with LFS.

## Author contributions

Concept and design: S.J. Shin, G. Peng, W. Wang

Development of methodology: S.J. Shin, Y. Yuan, W. Wang

Acquisition of data: J. Bojadzieva, L.C. Strong, P.L. Mai, S.A. Savage, M.L. Ballinger, D.M. Thomas

Analysis and interpretation of data: S.J. Shin, E.B. Dodd, Jingxiao Chen, G. Peng, C. Amos, W. Wang

Writing, review and/or revision of the manuscript: S.J. Shin, E.B. Dodd, J. Bojadzieva, C. Amos, S. A.

Savage, P. L. Mai, D. M. Thomas, M.L. Ballinger, L.C. Strong, W. Wang

Administrative, technical, or material support: E.B. Dodd

Study Supervision: W. Wang

