## Supplemental Figures for "Risk of differential cancer types over age in families with Li-Fraumeni syndrome: a validation study using multi-center cohorts"

**Supplementary Figure 1** Histograms of Calendar Years of Date of Birth and Death for pediatric sarcoma cohort

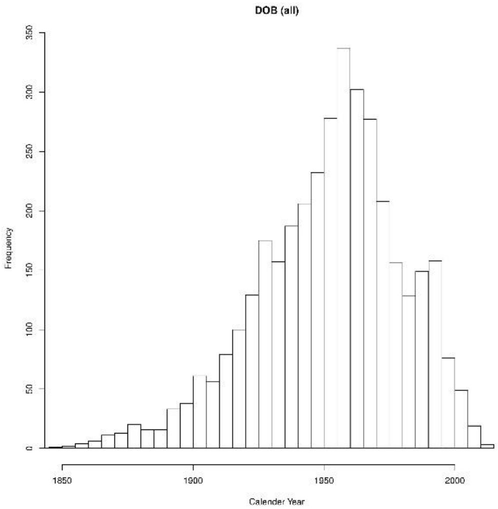


A. DOB

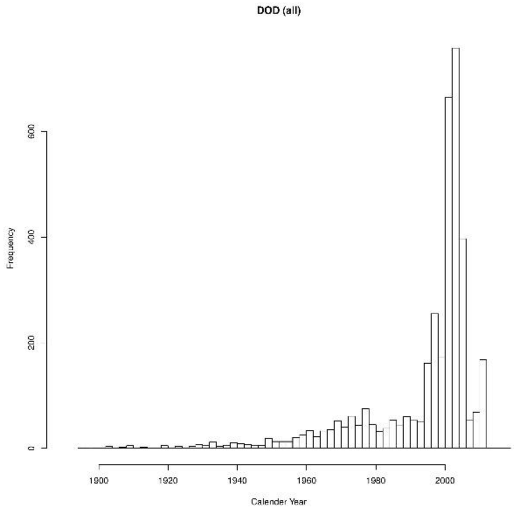


B. DOD

**Supplementary Figure 2** Mortality for US population.

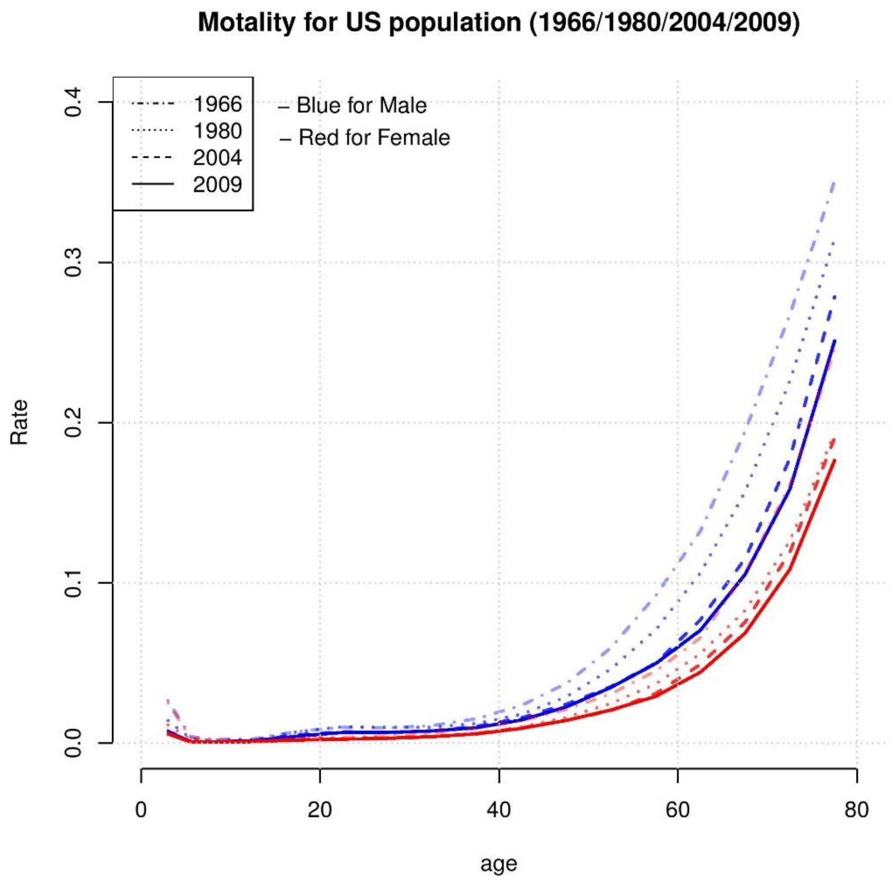
